# A Two-Heads-Bound State Drives KIF1A Superprocessivity

**DOI:** 10.1101/2025.01.14.632505

**Authors:** Lu Rao, Jan O. Wirth, Jessica Matthias, Arne Gennerich

**Affiliations:** Department of Biochemistry and Gruss Lipper Biophotonics Center, Albert Einstein College of Medicine, Bronx, NY 10461; Abberior Instruments America, Bethesda, Maryland 20814

## Abstract

KIF1A, a neuron-specific Kinesin-3 motor, is indispensable for long-distance axonal transport and nuclear migration, processes vital for neuronal function. Using MINFLUX tracking, we reveal that KIF1A predominantly adopts a two-heads-bound state, even under ATP-limiting conditions, challenging prior models proposing a one-head-bound rate-limiting step. This two-heads-bound conformation, stabilized by interactions between the positively charged K-loop and negatively charged tubulin tails, enhances microtubule affinity and minimizes detachment. The shorter neck linker facilitates inter-head tension, keeping the heads out of phase and enabling highly coordinated stepping. In contrast, Kinesin-1 (KIF5B) transitions to a one-head-bound state under similar conditions, limiting its processivity. Perturbing KIF1A’s mechanochemical cycle by prolonging its one-head-bound state significantly reduces processivity, underscoring the critical role of the two-heads-bound state in motility. These findings establish a mechanistic framework for understanding KIF1A’s adaptations for neuronal transport and dysfunction in neurological diseases.

## Introduction

Intracellular transport is essential for maintaining cellular organization and function, especially in neurons, where the efficient delivery of cargo over long distances is crucial for neuronal function and synaptic communication. Kinesin motor proteins play a central role in this process, with KIF1A, a member of the Kinesin-3 family, being a key driver in the anterograde transport of synaptic vesicle precursors, dense core vesicles, and other vital cargo along microtubules (MTs). Mutations in KIF1A cause severe neurodevelopmental and neurodegenerative conditions collectively referred to as KIF1A-associated neurological disorders (KAND). KIF1A is distinguished by its extraordinary processivity, often referred to as “superprocessivity,” enabling it to travel micrometer distances along MTs without detachment—a feature critical for its function in elongated neuronal axons.

KIF1A’s remarkable processivity is attributed to unique structural and mechanistic features that distinguish it from other kinesins. Key among these is its class-specific loop 12, known as the “K-loop,” which contains a KNKKKKK sequence of six positively charged lysine residues. This K-loop interacts with negatively charged α- and β-tubulin tails, enhancing MT binding. Additionally, KIF1A has a neck linker that is ∼3Å shorter than that of Kinesin-1 KIF5B, the founding member of the kinesin superfamily. We hypothesize that this shorter neck linker not only drives the ATP-dependent forward movement of KIF1A’s trailing motor domain but also helps establish sufficient tension between the two heads to keep them “out of phase”. This mechanism ensures that the leading motor domain remains attached to the MT while the trailing motor domain weakly binds and is pulled forward through neck-linker docking. Supporting this hypothesis, increasing KIF1A’s neck linker length by 3Å, by replacing its proline (P) with leucine (L) from KIF5B at the same position, significantly reduces run length^1^, suggesting impaired head-head coordination.

Effective tension transmission between the motor domains requires a sufficiently long-lived two-heads-bound state. However, prior studies suggested that KIF1A’s rate-limiting step involves a one-head-bound state^2,3^, which could increase the risk of detachment. To determine how KIF1A achieves highly processive motion despite this potential limitation, we leveraged the exceptional spatiotemporal resolution of the MINFLUX, a single-molecule localization technique that determines fluorophore positions by relating the zero intensity position of a donut-shaped excitation beam to the fluorophore’s unknown location^4–6^. This approach enabled precise tracking of KIF1A’s motor domains (**Fig. 1A**). Our findings reveal that, in contrast to KIF5B, KIF1A predominantly adopts a two-heads-bound state, even under limiting ATP conditions. This configuration, stabilized by the K-loop, facilitates enhanced inter-head tension via the shorter neck linker, maintaining the heads “out of phase” and reducing detachment risk. These findings provide a molecular basis for KIF1A’s superprocessivity and underscore its critical role in neuronal cargo transport.

**Figure 1.**
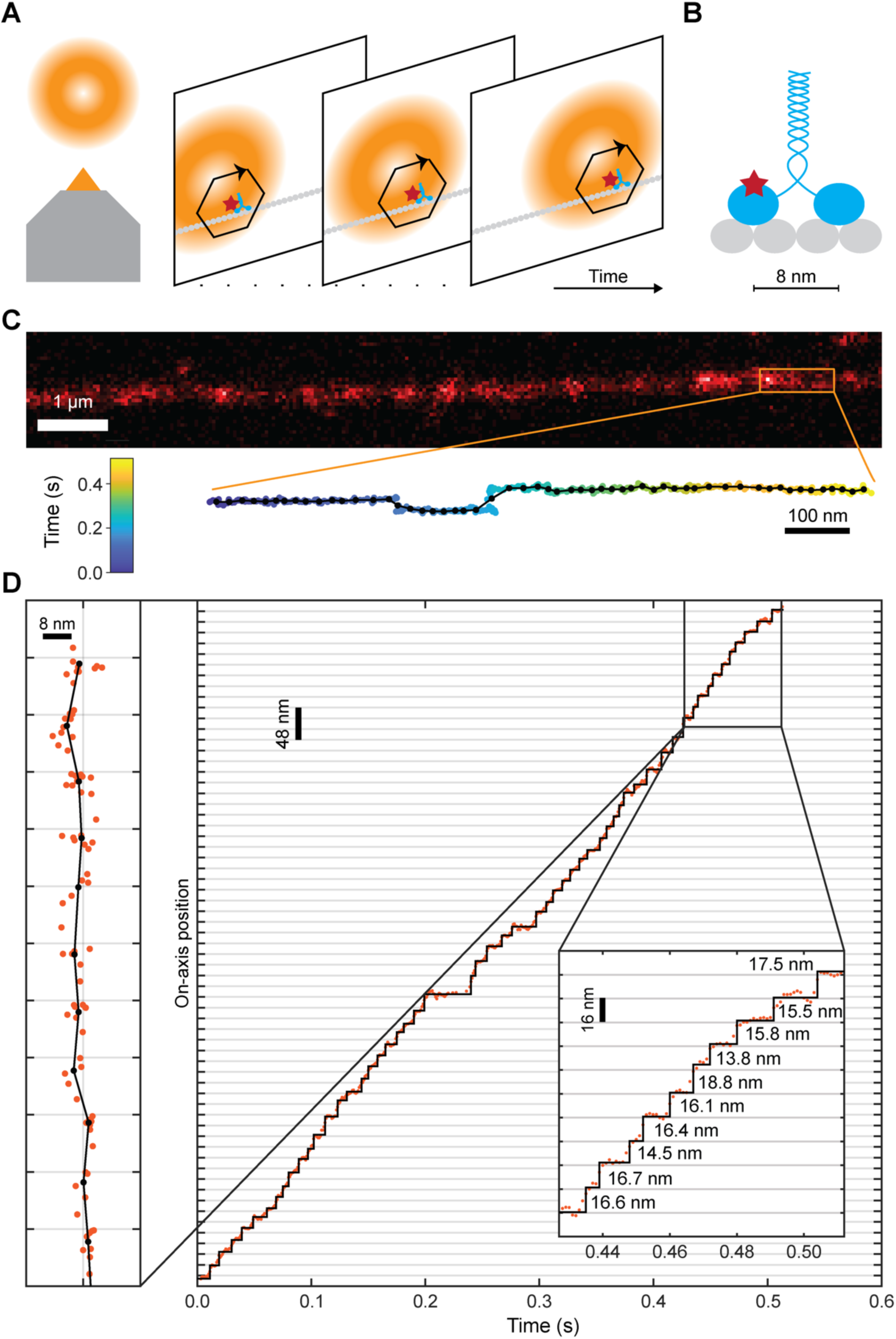
MINFLUX tracking of KIF1A. **(A)** Schematic of the MINFLUX tracking process, featuring a donut-shaped excitation beam. For each timepoint, the *xy*-position of a fluorophore (red star) is triangulated by probing a hexagonal pattern. During tracking, the beam follows the fluorophore, revealing the position of the motor protein (blue) along the microtubule (gray). **(B)** Schematic of the ybbR-tagged KIF1A construct (blue) used in this study, positioned on a microtubule filament (gray). The fluorophore (red start) is located approximately at the top-middle of the motor domain. **(C)** Confocal image of a single microtubule with a zoom-in view of a single track of ybbR-tagged KIF1A. The binned localizations (1 ms binning, color-coded by time) are overlaid with a fitted step trace (black dots and line). **(D)** On-axis position of the motor along the microtubule over time (orange dots), overlaid with the fitted stepwise motion (black line). The zoom-in at approximately 0.3 s shows the binned *xy*-coordinates overlaid with the fitted steps (left) and the measured step sizes (inlay). Pixel size in C: 50 nm.

## Results

### KIF1A Adopts a Two-Heads-Bound State at Both Physiological and Limiting ATP Levels

To investigate whether KIF1A exhibits a sufficiently long-lived two-heads-bound state under physiological conditions to enable the development of inter-head tension^1^, or if a one-head-bound state dominates KIF1A’s mechanochemical cycle as suggested by prior studies^2,3^, we measured the dynamics of KIF1A’s motor domains at a saturating ATP concentrations (2 mM). Using MINFLUX tracking, which provides ∼2 nm spatial precision and sub-millisecond temporal resolution, we resolved individual motor steps, even at KIF1A’s high velocity (2.3 µm/s, **Figs. 1&2B**).

To achieve high-resolution tracking, we inserted an 11-amino acid ybbR-tag^7^ at position 134-135 within KIF1A’s motor domain for site-specific labeling with the bright and photostable organic fluorophore LD655 (Lumidyne Technologies) (**Fig. 2A**). A tail-truncated, dimerized KIF1A construct (KIF1A (1–393)-LZ)^8^ with a leucine zipper was expressed to facilitate dimerization, and motor domains were sparsely labeled, ensuring that only one of the two carried an LD655 fluorophore (**Fig. 2A**). Sparse labeling was confirmed via photobleaching assays (**Suppl. Fig. 1**). Purified motors were introduced into a flow chamber containing taxol-stabilized MTs immobilized on coverslips. The ybbR-tagged motors showed velocities and run lengths comparable to the WT motor (**Fig. 2B**), validating their functional integrity. To reliably visualize the motor’s one-head-bound state, we used 2D-MINFLUX tracking with a focus on spatial precision over temporal resolution. This approach allowed us to recorded KIF1A motility along individual MTs (**Fig. 1C**) and analyzed stepwise displacements using a custom MATLAB algorithm (**Fig. 1D**). Despite KIF1A’s rapid velocity at saturating ATP (2.3 µm/s), the detected motion predominantly consisted of ∼16 nm steps (**Fig. 1D**, inset). The step-size histograms confirmed this ∼16 nm peak, with occasional larger steps likely resulting from missed transitions due to the motor’s speed (**Fig. 3C**, right). These findings indicate that KIF1A predominantly maintains a two-heads-bound state under physiological conditions.

**Figure 2.**
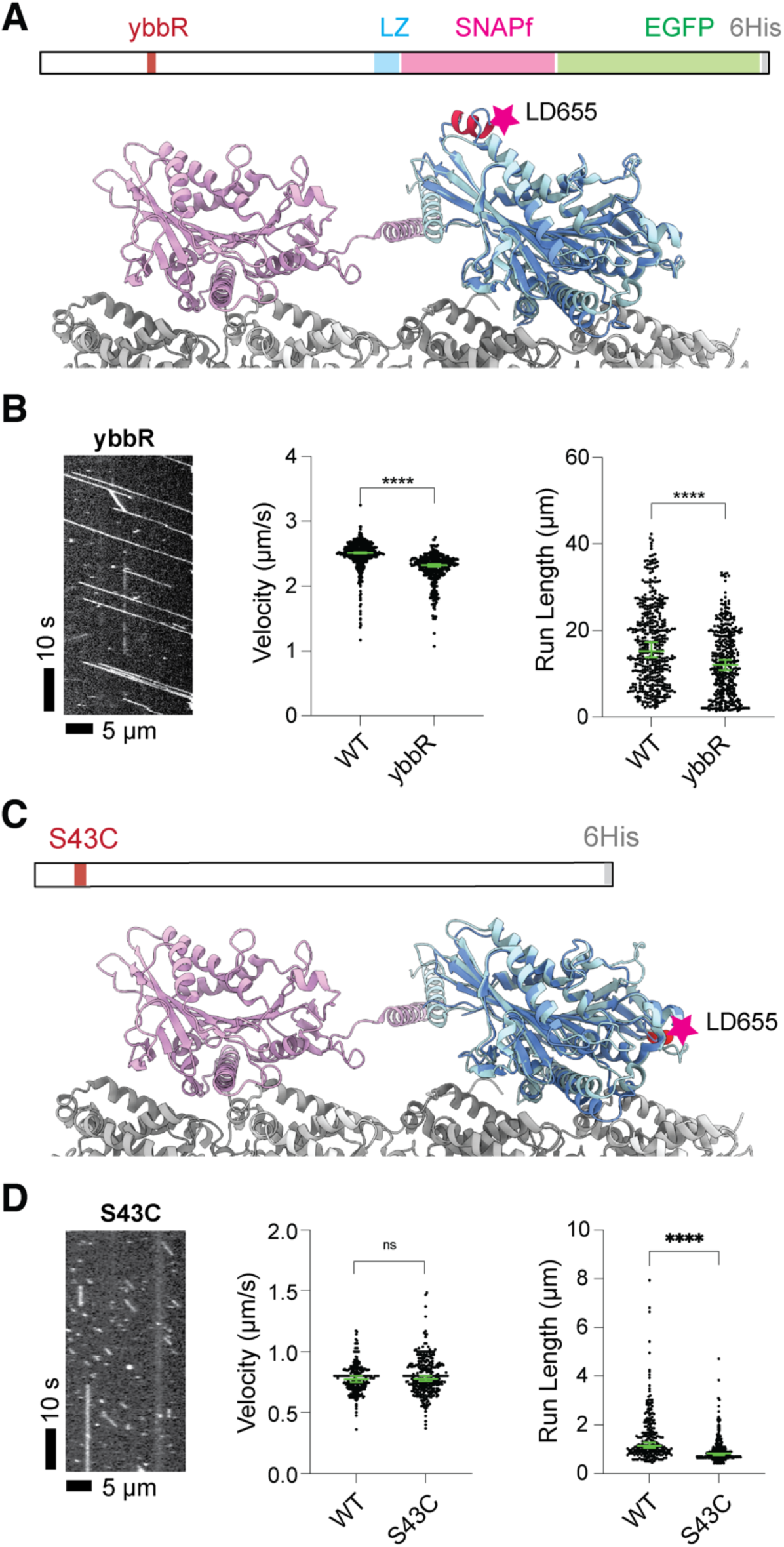
KIF1A and KIF5B constructs for MINFLUX tracking exhibit similar behavior to WT. **(A)** (Top) Schematic representation of the KIF1A construct for MINFLUX tracking. A ybbR-tag^7^ was inserted into the motor domain of KIF1A, as previously described^1^. Dimerization of KIF1A was facilitated by a leucine zipper (LZ), and the ybbR-tag was labeled with CoA-LD655 at low labeling ratio. (Bottom) ColabFold-predicted structure^72^ of the ybbR-tagged KIF1A motor domain overlaid with the trailing head from a cryo-EM structure of KIF1A bound to microtubules (PDB 8UTN)^1^. The labeling position was indicated with a red star. **(B)** Kymograph of the ybbR-tagged KIF1A construct, showing processive movement along microtubules with slightly reduced velocity and run length compared to WT. WT: KIF1A(1–393)-LZ-SNAPf-EGFP-6His (**Suppl. Fig. 5**), labeled with SNAP-TMR; ybbR: KIF1A(1–393, ybbR inserted between 134 and 135)-LZ-SNAPf-EGFP-6His, labeled with CoA-LD655. Green bars in the velocity and run length graphs indicated the median with 95% confidence interval (CI). WT: n=419; ybbR: n=371. Velocity: WT: 2.51 [2.50, 2.52] µm/s; ybbR: 2.32 [2.30, 2.34] µm/s; **** *P*<0.0001 (unpaired *t*-test with Welch’s correction). Run length: WT: 15.26 [13.71, 17.26] µm; ybbR: 12.10 [10.90, 13.19] µm; **** *P*<0.0001 (Kolmogorov-Smirnov test). **(C)** (Top) Schematic representation of the KIF5B construct used for MINFLUX tracking. A single mutation (S43C) was introduced into the motor domain of a cysteine-light mutant (clm) KIF5B construct^67^. The Cys43 residue was labeled with maleimide-LD655 at low labeling ratio. (Bottom) ColabFold-predicted clm-KIF5B(S43C) motor domain overlaid with the trailing head from a cryo-EM structure of KIF1A bound to microtubules (PDB 8UTN)^1^. The labeling position is indicated with a red star. **(D)** Kymograph of the S43C KIF5B construct, demonstrating processive movement along microtubules with similar velocity and slightly decreased run length compared to WT. WT: KIF5B(1–490)-SNAPf-EGFP-6His, labeled with SNAP-TMR; S43C: KIF5B(clm, S43C)-6His, labeled with maleimide-LD655. Green bars in the velocity and run length graphs indicate the median with 95% CI. WT: n=254; S43C: n=303. Velocity: WT: 0.77 [0.75, 0.80] µm/s; S43C: 0.78 [0.76, 0.80] µm/s; ns, *P*=0.1471 (unpaired *t*-test with Welch’s correction). Run length: WT: 1.13 [1.07, 1.28] µm; S43C: 0.84 [0.77, 0.84] µm; **** *P*<0.0001 (Kolmogorov-Smirnov test).

**Figure 3.**
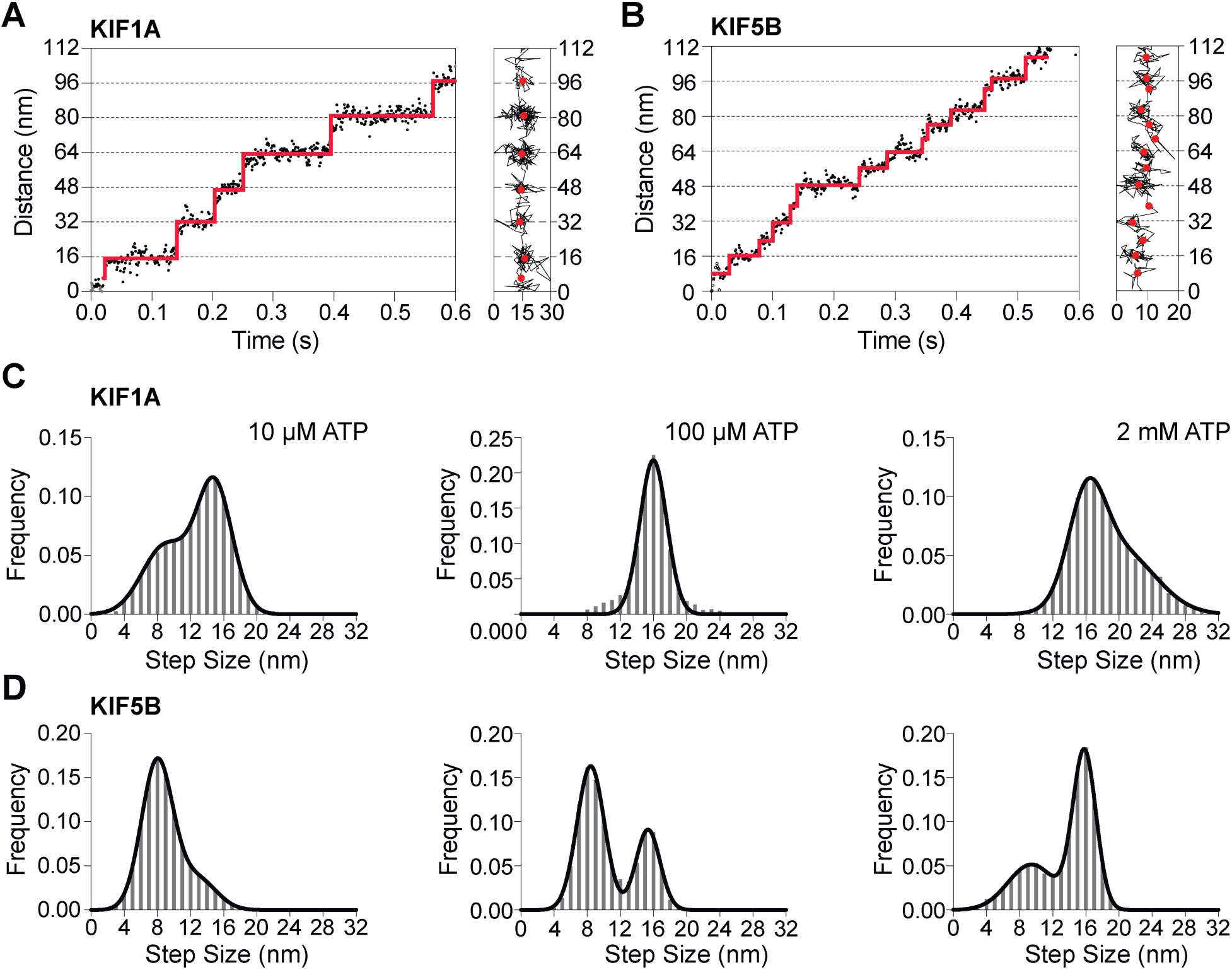
KIF1A exhibits a distinct stepping behavior compared to KIF5B. **(A and B)** Example traces of KIF1A and KIF5B stepping along microtubules at 10 µM ATP. (Left) 1D trace representing movement along the microtubule axis; (Right) 2D trace showing motor trajectories in the xy-plane. **(C)** Step size histograms of KIF1A at various ATP concentration. At 10 µM ATP, the histogram was fitted with a sum of two Gaussians, with the centers as 8.95 [8.67, 9.28] nm (mean with 95% CI) and 14.90 [14.78, 15.02] nm. At 100 µM ATP, the histogram was fitted with a single Gaussians, with the center at 15.98 [15.89, 16.07] nm. At 2 mM ATP, the histogram was fitted with the sum of two Gaussians, with the centers at 16.10 [16.01, 16.33] nm and 20.25 [19.63, 22.09] nm. **(D)** Step-size histograms of KIF5B at various ATP concentrations. All histograms were fitted with the sum of two Gaussians. At 10 µM ATP, the centers were 7.98 [7.93, 8.02] nm (mean with 95% CI) and 12.82 [12.45, 13.08] nm. At 100 µM ATP, the center was 8.42 [8.35, 8.50] nm. At 2 mM ATP, the centers were 9.45 [9.27, 9.66] nm and 15.83 [15.79, 15.86] nm.

We further examined KIF1A’s behavior under rate-limiting ATP conditions, hypothesizing that the trailing motor domain might hydrolyze ATP, release phosphate, and transition into a weak MT-binding ADP state, potentially dissociating before the leading domain binds ATP and undergoes neck-linker docking. At 100 µM ATP, where the velocity decreased from ∼2.3 µm/s to ∼720 nm/s (**Suppl. Fig. 2**), the step-size histogram still displayed a single peak at ∼16 nm (**Fig. 3C**, middle), suggesting that the larger steps observed at 2 mM ATP were due to missed transitions. At 10 µM ATP, where the velocity further declined to ∼270 nm/s (**Suppl. Fig. 2**), a population of ∼8 nm steps emerged, indicative of a sufficiently long-lived one-head-bound state, though the dominant peak remained at ∼16 nm (**Fig. 3C**, left). These findings demonstrate that even under rate-limiting conditions, KIF1A predominantly maintains a two-heads-bound state, with the trailing head entering a one-head-bound only briefly and under non-physiological conditions where ATP binding is significantly delayed.

### KIF5B Adopts a One-Head-Bound State Before ATP Binds to the Leading Head

A previous study suggested that Kinesin-1 (KIF5B), like KIF1A, spends most of its time in a two-heads-bound state at saturating ATP concentrations^9^. However, it was proposed that KIF5B transitions to a one-head-bound state while waiting for ATP^9^, a finding recently confirmed by a MINFLUX study^10^. Interestingly, earlier work indicated that KIF1A’s trailing head detaches more rapidly than that of KIF5B^2,3^, implying that KIF1A might favor a one-head-bound state. In contrast to KIF5B, our results reveal that KIF1A predominantly maintains a two-heads-bound state, even under rate-limiting ATP conditions (**Fig. 3C**). To exclude potential biases from experimental conditions, we directly compared the stepping behavior of KIF1A and KIF5B under identical conditions. For KIF5B, we used a cysteine-light mutant^11^ labeled with LD655 at residue 43 (S43C)^9^ in its motor domain to enable single-molecule tracking.

Given that KIF5B is significantly less processive than KIF1A (**Fig. 2A&B**), we hypothesized that KIF5B’s trailing head detaches more quickly, leaving the motor in a vulnerable one-head-bound state, which may account for its reduced processivity (**Fig. 2B&C**). To test this, we used 2D-MINFLUX tracking to record KIF5B’s motor dynamics and analyzed stepwise displacements. At physiological ATP concentrations, KIF5B’s step-size histogram displayed a major peak at ∼16 nm (**Fig. 3D**, right), similar to KIF1A (**Fig. 3C**, right). However, KIF5B also exhibited a notable fraction of ∼8 nm steps. As ATP concentrations decreased, the ∼8 nm peak became dominant (**Fig. 3D**, middle), and at 10 µM ATP, ∼16 nm steps were almost absent (**Fig. 3D**, left). At this low ATP concentration, KIF1A and KIF5B exhibited comparable walking speeds (**Suppl. Fig. 2**), suggesting similar ATP turnover rates. Despite this, the higher frequency of ∼8 nm steps for KIF5B indicates a prolonged one-head-bound state, consistent with a faster detachment rate of the trailing head. This likely explains KIF5B’s lower processivity compared to KIF1A.

### KIF1A’s Two-Heads-Bound State Arises from a K-Loop-Dependent Increased MT Affinity in the Motor’s ADP State

The long-lived two-heads-bound state of KIF1A observed in our experiments suggests that the trailing head remains MT bound even after ATP hydrolysis and phosphate release, persisting in the weak MT-binding ADP state. If true, KIF1A’s MT affinity in the ADP state should be significantly stronger than that of KIF5B, which predominantly adopts a one-head-bound state under rate-limiting ATP conditions (**Fig. 3D**, left), unlike KIF1A (**Fig. 3C**, left). To test this hypothesis, we used total internal reflection fluorescence (TIRF) microscopy to image SNAPf-tagged motors binding to MTs immobilized on a coverslip. While both KIF1A and KIF5B exhibited strong MT-binding in the nucleotide-free apo state (**Fig. 4A**), KIF1A showed significantly longer MT-binding durations compared to KIF5B in the presence of 1 mM ADP (**Fig. 4B**, WT KIF5B and WT KIF1A).

**Figure 4.**
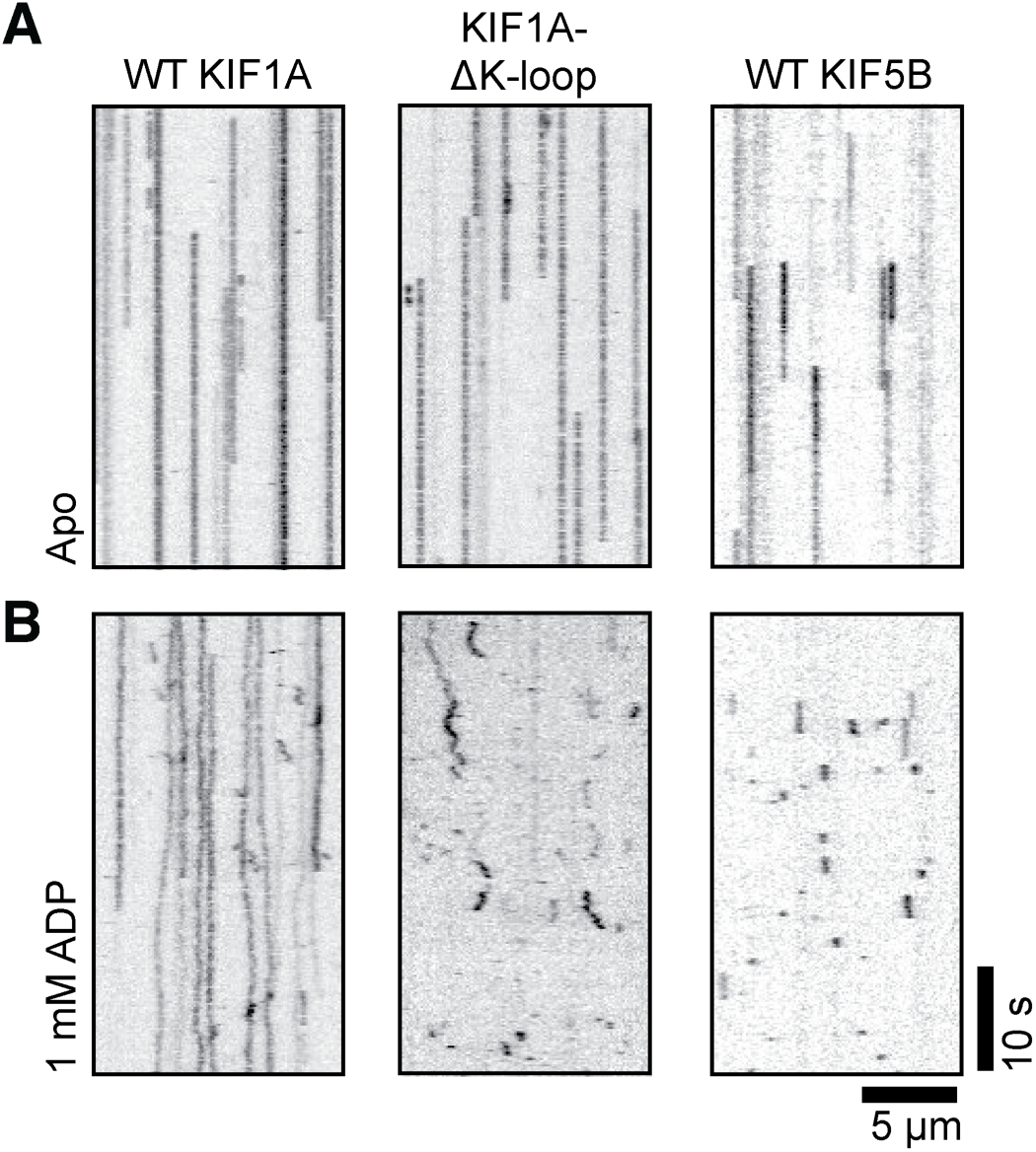
KIF1A exhibits prolonged dwell time on microtubules at ADP state. **(A)** Representative kymographs showing the binding of KIF1A, KIF1A with K-loop deletion, and KIF5B to microtubules in the apo state. **(B)** Representative kymographs showing the binding of KIF1A, KIF1A with K-loop deletion, and KIF5B to microtubules in the presence of 1 mM ADP.

To investigate whether KIF1A’s strong MT association in the ADP state depends on its K-loop, we tested a KIF1A construct where the K-loop was replaced with KIF5B’s corresponding L12-loop^8^. As expected, in the absence of the K-loop, KIF1A’s MT-binding duration was markedly reduced, resembling the brief interactions of KIF5B in the presence of 1 mM ADP (**Fig. 4B**). This supports the hypothesis that interactions between the positively charged K-loop and the negatively charged α- and β-tubulin tails^1^ stabilize KIF1A’s two-heads-bound state, even when the trailing head is in the ADP-bound state, in contrast to KIF5B.

Although KIF1A binds less tightly to MTs in the presence of 1 mM ADP compared to nucleotide-free (apo) conditions—exhibiting some diffusion along the MT—it retains similar overall MT-bound times (**Fig. 4A&B**). This contrasts with a previous study suggesting that KIF1A’s MT binding in the presence of 1 mM ADP is five times shorter than in apo conditions^2^. To resolve this discrepancy, we replicated the conditions of the earlier study, which used a kinesin chimera dimerizing KIF1A motor domains via the *D. melanogaster* kinesin-1 neck coil (NC)^2^. Consistent with prior findings^2^, when we fused our human KIF1A construct to the human KIF5B NC (KIF1A-KIF5B NC), the resulting chimera exhibited strong MT binding in the apo state, similar to wild-type KIF1A, but showed only brief MT interactions in the presence of ADP (**Suppl. Fig. 3**). This result suggests that replacing KIF1A’s native NC with KIF5B’s diminishes its MT affinity in the ADP state. We conclude that prior kinetic studies using chimeric constructs may not fully represent KIF1A’s behavior, highlighting the importance of retaining KIF1A’s native structural features for accurately characterizing its motility properties.

### KIF1A Exhibits a Shorter One-Head-Bound State Compared to KIF5B

To further investigate the stepping mechanism of KIF1A, we analyzed the dwell times between steps and compared them to KIF5B. Since the dwell times between consecutive ∼16 nm steps correspond to two full hydrolysis cycles and primarily reflect walking speed, our analysis focused on the shorter dwell times between consecutive ∼8 nm steps (**Fig. 5A**), which were most prominent at 10 µM ATP. These shorter dwell times represent the duration of the one-head-bound state and the subsequent ATP-driven transition.

**Figure 5.**
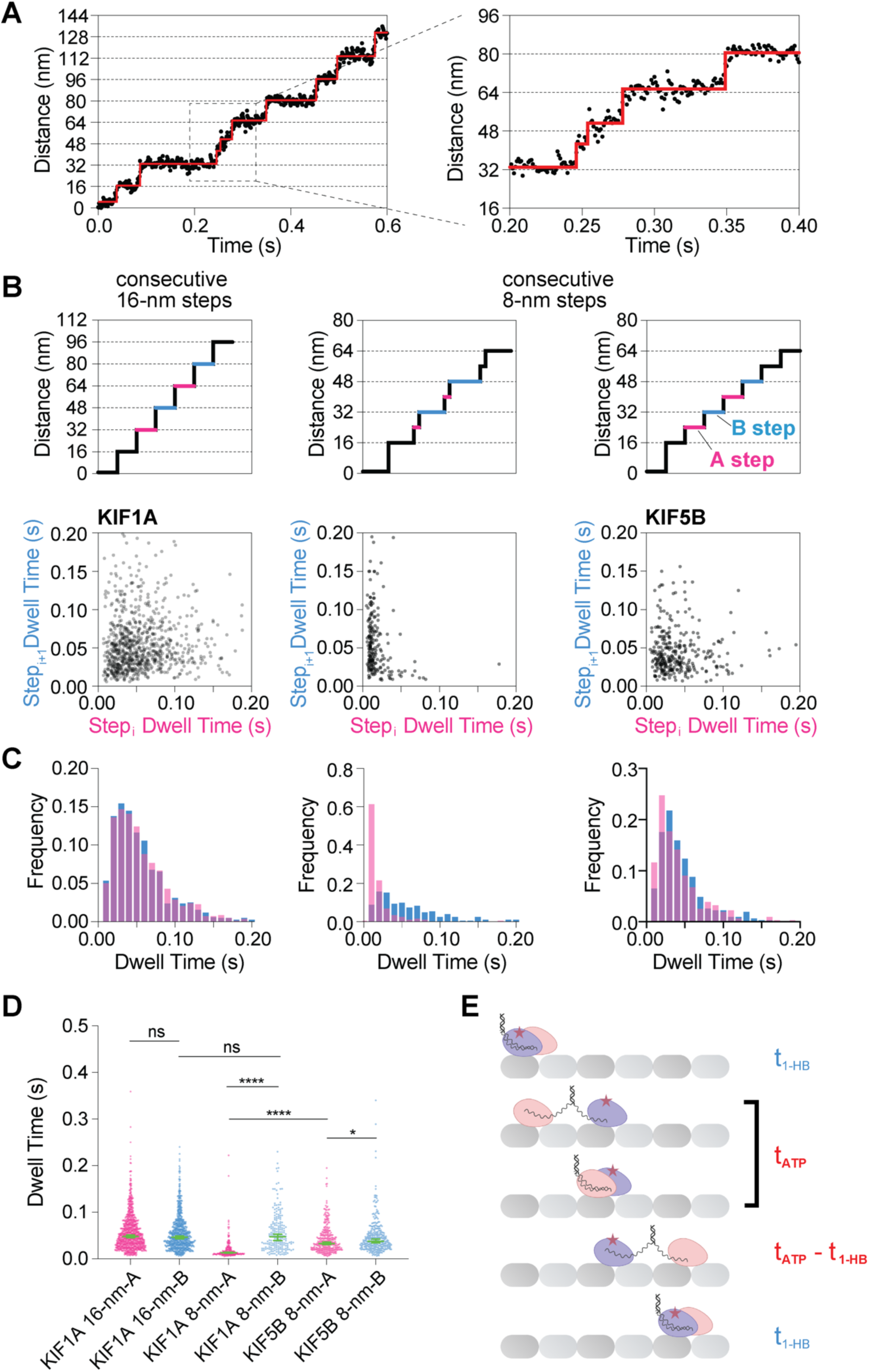
KIF1A’s sub-steps exhibit shorter dwell times compared to KIF5B. **(A)** Example trace of KIF1A’s sub-steps, with zoom-in showing the short dwell time of sub-steps. **(B)** (Top) Illustrations of the dwell time analysis. Consecutive steps of either 16 ± 2 nm or 8 ± 2 nm were detected following a 16-nm step. Dwell times of A-steps (red) and B-steps (blue) were analyzed separately and plotted as correlation graphs (Bottom). KIF1A 16-nm: n=847, Pearson *r*=0.05060 [−0.01682, 0.1176] (95% CI), *P*=0.1412; KIF1A 8-nm: n=205, Pearson *r*=-0.2283 [−0.3542, - 0.09418], *P*=0.0010; KIF5B 8-nm: n=312, Pearson *r*=0.008229 [−0.1029, 0.1192], *P*=0.8849. **(C)** Histograms of the dwell times, showing that KIF1A has shorter intermediate states than KIF5B. Colors correspond to (*B*). **(D)** Scatter plots of the dwell times, with the green bars indicating medians with 95% CI. KIF1A 16-nm-A: 0.048 [0.045, 0051] s; KIF1A 16-nm-B: 0.046 [0.043, 0.049] s; KIF1A 8-nm-A: 0.013 [0.012, 0.014] s; KIF1A 8-nm-B: 0.047 [0.039, 0.053] s; KIF5B 8-nm-A: 0.032 [0.030, 0.036] s; KIF5B 8-nm-B: 0.038 [0.034, 0.042] s. Kolmogorov-Smirnov test was used for the data sets. ns, *P*=0.5412 (KIF1A 16-nm-A and -B), *P*=0.6458 (KIF1A 16-nm-B and KIF1A 8-nm-B); *, *P*=0.0119; ****, *P*<0.0001. **(E)** Illustrations of the dwell times of a labeled motor domain (purple motor domain with a red star indicating LD655) in different states. The motor alternates between detachment (blue labels) and attachment (red labels) states. During detachment, the motor is in a one-head-bound state (t_1-HB_). When attached, it undergoes ATP-waiting and ATP-hydrolysis (t_ATP_), and either detaches or remains bound until the leading head binds ATP (t_ATP_ – t_1-HB_).

To accurately detect these distinct stepping patterns, we analyzed the data using two criteria: (1) Consecutive ∼16 nm steps were identified based on a step size of 16 ± 2 nm (**Fig. 5B**, top left) and were alternatingly assigned as A-steps (red) and B-steps (blue; **Fig. 5B**); (2) For ∼8 nm steps, three conditions were applied to ensure a clear separation of alternating A- and B-steps (Fig. 5B, top middle and right). First, the initial step in a series of steps must measure 16 ± 2 nm. Second, subsequent steps within the series must fall within 8 ± 2 nm. Notably, the ∼8 nm criterion was chosen because the sub-step sizes for both KIF1A and KIF5B are centered around ∼8 nm at 10 µM ATP (Fig. 3C, D). Removing this criterion yielded similar conclusions (**Suppl. Fig. 4**). These findings further support that KIF1A spends most of its time in a two-heads-bound state and only briefly transitions to a one-head-bound state. For ∼8 nm steps, the A-step represents the trailing head detaching from the MT and the motor entering a one-head-bound state, while the B-step corresponds to the unbound head reaching forward and binding to the MT. In the case of ∼16 nm steps, the A- and B-steps simply reflect consecutive motor steps.

By plotting the dwell times following A-steps (t_A_) against those following B-steps (t_B_), we identified distinct distributions for the different stepping patterns and constructs (**Fig. 5B**, bottom). For KIF1A’s ∼16 nm steps, the dwell times showed no correlation, as expected for independent ATP-binding and hydrolysis events (**Fig. 5B**, bottom left). In contrast, its ∼8 nm steps displayed a strong correlation, with short dwell time after the A-step followed by a long dwell time after the B-step (**Fig. 5B**, bottom middle). The long dwell times were similar to those observed between consecutive ∼16 nm steps (**Fig. 5 C&D**), supporting our identification of the A-step as the motor entering the one-head-bound state.

In contrast, the dwell times after KIF5B’s ∼8 nm A- and B-steps showed no correlation (**Fig. 5B**, bottom right). Comparing the dwell times after the A-step between KIF1A and KIF5B highlights the reduced duration of the one-head-bound state for KIF1A (**Fig. 5 C&D**). Interestingly, for KIF5B, the dwell time following the ∼8 nm B-step is only slightly longer than that of the A-step. This suggests that KIF5B’s one-head-bound time (t_A_ = t_1-HB_) is comparable to the time required for the leading head to bind ATP and hydrolyze it (t_B_ = [t_ATP_ + (t_ATP_ – t_1-HB_)]) (**Fig. 5E**). Consequently, t_1-HB_ closely approaches t_ATP_, indicating that KIF5B’s trailing head detaches almost immediately after the leading head binds to a new forward site. In contrast, KIF1A’s trailing head remains closely associated with the MT until the leading head binds ATP, ensuring a longer two-heads-bound state.

### Increasing KIF1A’s One-Head-Bound Time Leads to Decreased Processivity

To investigate how KIF1A’s trailing head steps around the leading head, we initially engineered a KIF1A heterodimer using a heterodimerizing leucine zipper^12,13^ (Fig. 6A). In this construct, residue K41 was replaced with the unnatural amino acid 4-Azido-L-phenylalanine (AzF) via amber codon suppression. This substitution allowed for site-specific labeling with DBCO-LD655 through click chemistry (**Fig. 6A**). Control experiments showed that the heterodimer without amber codon dimerized efficiently and exhibited processivity comparable to the WT homodimer (**Suppl. Fig. 5A-C**). However, introducing K41F significantly reduced KIF1A’s processivity, even though the position is analogous to KIF5B’s S43C (**Fig. 6B**).

**Figure 6.**
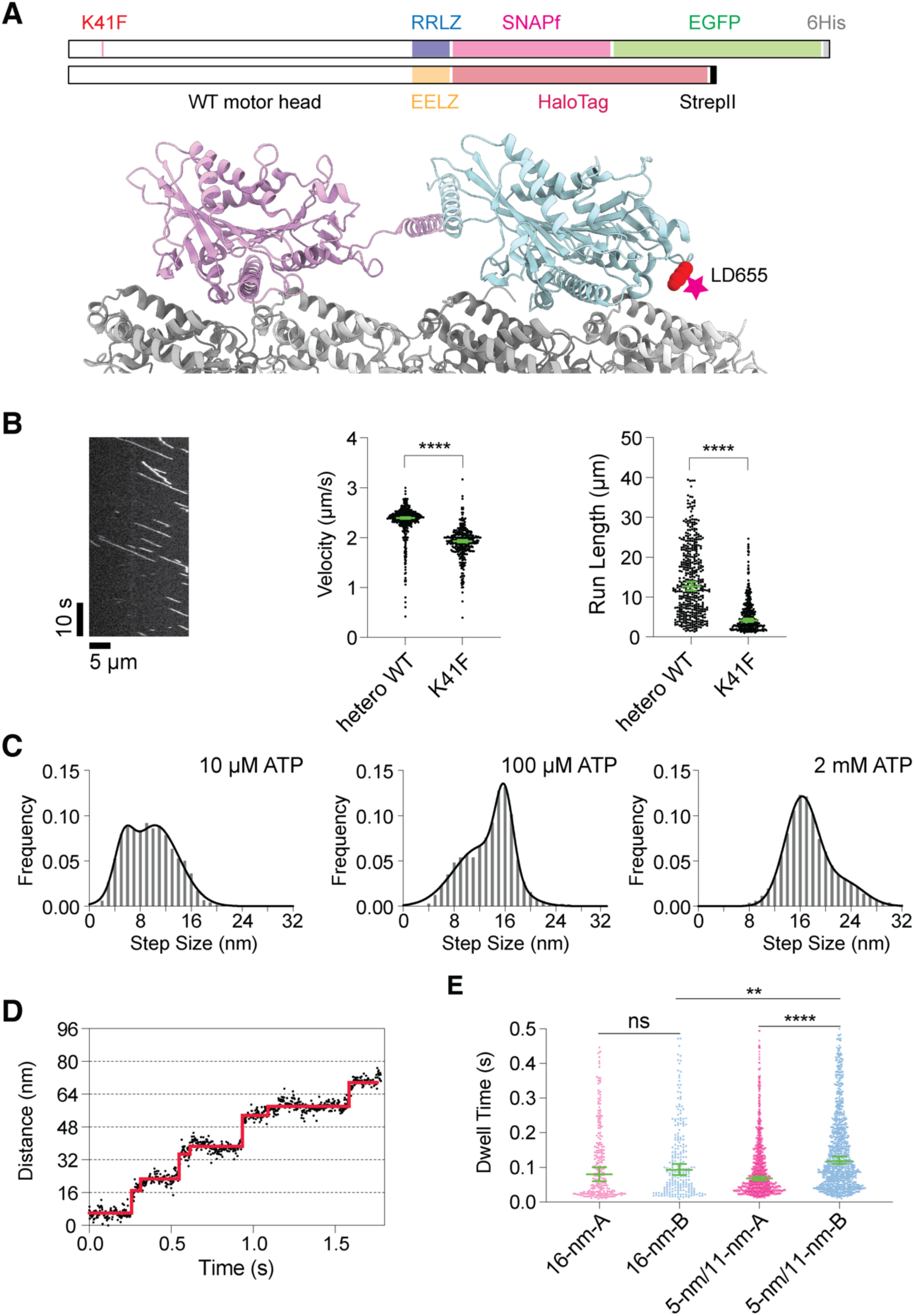
KIF1A K41F-LD655 shows increased one-head-bound state time and reduced run length. **(A)** (Top) Schematic representation of the KIF1A heterodimeric constructs with an unnatural amino acid inserted at K41 and labeled with LD655 for MINFLUX tracking. The constructs were dimerized via a leuzine-zipper-like coiled coil with opposite charges (RRLZ and EELZ). The RRLZ monomer bears the unnatural amino acid and a C-terminal SNAPf-EGFP-6His tag, while the EELZ monomer has a C-terminal HaloTag-StrepII. (Bottom) The red sphere represents K41 on the structure of KIF1A bound to microtubules (PDB 8UTN), indicating the labeling position. **(B)** Kymograph showing the motility of the labeled KIF1A K41F construct. Heterodimeric WT was labeled with SNAP-TMR. Green bars in the velocity and run length graphs indicate the median with 95% CI. Velocity: heterodimeric WT: 2.39 [2.38, 2.41] µm/s, n=440; K41F: 1.93 [1.91, 1.96] µm/s, n=334. Run length: heterodimeric WT: 12.70 [11.66, 13.84] µm; K41F: 4.16 [3.81, 4.58] µm. **(C)** Step-size histograms of KIF1A K41F at various ATP concentration, fitted with the sum of two Gaussians. 10 µM ATP: 5.34 [5.14, 5.62] nm (mean with 95% CI) and 10.34 [9.96, 10.83] nm. 100 µM ATP: 12.16 [11.31, 12.65] nm. 2 mM ATP: 16.23 [15.84, 16.39] nm and 23.02 [17.80, 23.80] nm. **(D)** Example trace of KIF1A K41F stepping at 10 µM ATP. **(E)** Scatter plots of the dwell times of KIF1A K41F. Color corresponds to Figure 5. Green bars indicate the median with 95% CI. 16-nm-A: n=244, 0.080 [0.060, 0.100] s; 16-nm-B: 0.093 [0.078, 0.110] s; 5-nm/11-nm-A: n=932, 0.068 [0.062, 0.074] s; 5-nm/11-nm-B: 0.118 [0.110, 0.132] s.

Using MINFLUX tracking, we analyzed the motility of the labeled construct. At 2 mM ATP, the K41F heterodimer predominantly exhibited ∼16-nm steps, similar to the ybbR-tagged WT KIF1A (**Fig. 6C**, right). At 100 µM ATP, smaller sub-steps became evident (**Fig. 6C**, middle), and at 10 µM ATP, the motor predominantly displayed ∼5 nm and ∼11 nm steps (**Fig. 6C**, left, and **Fig. 6D**). Analysis of the dwell times for these sub-steps revealed that the one-head-bound state in the K41F construct persisted five times longer than in the ybbR-tagged KIF1A (**Fig. 6E**). These findings demonstrate that reducing the time KIF1A spends in the two-heads-bound state and augmenting the one-head-bound state leads to reduction of the processivity of KIF1A. This result highlights the critical role of the two-heads-bound state in maintaining KIF1A’s efficient motility.

## Discussion

Efficient intracellular transport is critical for neuronal function, where long axons demand robust and processive motor proteins to transport essential cargo. Our study reveals key molecular mechanisms underlying KIF1A’s remarkable superprocessivity, distinguishing it from other kinesin motors, particularly Kinesin-1 (KIF5B). Using MINFLUX tracking, we demonstrate that KIF1A predominantly adopts a two-heads-bound state, even under ATP-limiting conditions. This behavior, coupled with its unique structural features such as the positively charged K-loop^3,14^ and shorter neck linker^1^, enables KIF1A to maintain inter-head tension, prevent premature detachment, and sustain highly processive motion. These findings provide critical insights into how KIF1A achieves efficient transport over long distances, a necessity for neuronal health and function.

### KIF1A’s Two-Heads-Bound State Enhances Processivity

Our analysis of dwell times after the A-step (t_1-HB_) and after the B-step [t_ATP_ + (t_ATP_ – t_1-HB_)] reveals a stark contrast between KIF5B and KIF1A. For KIF5B, t_1-HB_ ≈ t_ATP_ + (t_ATP_ – t_1-HB_) and thus t_1-HB_ ≈ t_ATP_, indicating that the motor spends the majority of its mechanochemical cycle in the one-head-bound state while waiting for ATP^10^. In contrast, for KIF1A, t_1-HB_ << t_ATP_ + (t_ATP_ – t_1-HB_), resulting in t_1-HB_ << t_ATP_. This highlights that KIF1A predominantly occupies a two-heads-bound state during its mechanochemical cycle.

This findings challenge prior models suggesting that KIF1A spends most of its cycle in a one-head-bound state^2,3^. Instead, our results demonstrate that KIF1A maintains a two-heads-bound state under both physiological and rate-limiting ATP conditions. This longer-lived two-heads-bound state facilitates inter-head tension generation via KIF1A’s the shorter neck linker^1^, ensuring that the motor domains remain out of phase and reducing the likelihood of detachment. This mechanism allows KIF1A to traverse MTs over micrometer distances^15,16^, a critical feature for efficient cargo transport in neuronal axons^17^.

We further hypothesize that ATP binding in the leading head triggers partial neck-linker docking, facilitating the detachment of the trailing ADP-bound head and transitioning the motor into a one-head-bound state—a process termed “ATP-triggered detachment”^18^. In contrast, Kinesin-1 (KIF5B), which is more prone to adopting a one-head-bound state, may not require neck-linker docking for this transition. Some studies suggest that in KIF5B, the detachment of the trailing head might even precede ATP binding in the leading head^19,20^, following a “front-head gating” mechanism^18^. However, whether ATP binding in KIF1A induces partial neck-linker docking and hydrolysis drives full docking, as in Kinesin-1—where ATP hydrolysis is required for full neck-linker docking^21^—remains an open question.

### Role of the K-Loop in Stabilizing the Two-Heads-Bound State

Our results reveal that KIF1A’s positively charged K-loop plays a pivotal role in stabilizing its two-heads-bound state. The K-loop interacts with the negatively charged α- and β-tubulin tails^1^, enhancing MT-binding affinity. This interaction is particularly crucial when the trailing head transitions to the weak MT-binding ADP state, ensuring that KIF1A maintains its two-heads-bound configuration. In contrast, KIF5B lacks this K-loop, and its weaker MT affinity in the ADP state results in a higher frequency of trailing head detachment.

Replacing KIF1A’s K-loop with KIF5B’s L12-loop significantly reduced MT-binding duration in the ADP state, further corroborating the importance of the K-loop. This structural feature allows KIF1A to remain closely associated with MTs even under conditions where ATP binding is delayed, providing a molecular basis for its superior processivity compared to KIF5B.

### One-Head-Bound State as a Limiting Factor for KIF1A Processivity

Our engineered KIF1A construct with a K41F substitution illustrates the impact of increasing the one-head-bound time on processivity. While the construct retained its ability to perform ∼16 nm steps at saturating ATP, smaller sub-steps emerged at lower ATP concentrations. At 10 µM ATP, the motor primarily exhibited ∼5 nm and ∼11 nm sub-steps, with a five-fold increase in the dwell time of the one-head-bound state. As a result, the K41F construct exhibited significantly reduced processivity compared to the wild-type motor, highlighting the critical role of the two-heads-bound state in ensuring efficient cargo transport.

The duration of the one-head-bound state depends on two key factors: (1) the time required to transition from the two-heads-bound state and (2) the on-rate of the tethered ADP-bound head binding to a new MT site. We hypothesize that the K-loop enhances KIF1A’s superprocessivity by stabilizing the trailing head’s interactions with the MT when ADP-bound, allowing sufficient time for interhead tension to develop, and by accelerating the on-rate of the tethered ADP-bound head, enabling rapid rebinding to the MT.

In the canonical Kinesin-1 stepping cycle, ATP hydrolysis in the leading MT-bound head induces full neck-linker docking^21^, propelling the ADP-bound tethered head forward. At this critical point, the tethered head either rapidly binds the MT and releases ADP or the post-hydrolysis head detaches before the tethered head binds, resulting in loss of processivity. For KIF1A, we propose the K-loop stabilizes the two-heads-bound state and accelerates tethered head rebinding, ensuring efficient head-head coordination, inter-head tension, and processive motion, even under ATP-limiting conditions.

### Implications for Neuronal Transport and Disease

Our findings underscore the unique adaptations of KIF1A for long-distance cargo transport in neurons. The combination of its shorter neck linker^1^, positively charged K-loop^3,14^, and ability to sustain a two-heads-bound state, even under ATP-limiting conditions, distinguishes KIF1A as a highly processive motor. In contrast, KIF5B’s higher propensity for transitioning to a one-head-bound state explains its lower processivity and limited suitability for long-range transport tasks. These insights have broad implications for understanding kinesin motor function in health and disease. Mutations in KIF1A, linked to KIF1A-associated neurological disorders (KAND)^17^, often impair motor function and disrupt cargo transport, leading to severe neurodevelopmental and neurodegenerative conditions^17,22–65^. By elucidating the molecular basis of KIF1A’s superprocessivity, our findings provide a foundation for developing therapeutic strategies aimed at restoring motor function in KAND patients^66^.

## Materials and Methods

### Plasmids and Constructs

The ybbR-tagged KIF1A construct was derived from a previously described KIF1A(1–393)-LZ-SNAPf-EGFP-6His construct^1^. The ybbR-tag was inserted using Q5 mutagenesis (New England Biolabs, # E0554S). KIF5B(1-490)-SNAPf-EGFP-6His and KIF1A ΔK-loop constructs were obtained from our previous study^1^. The cys-light (cl) KIF5B S43C mutant was a generously provided by the laboratory of Dr. Ronald D. Vale (University of California, San Francisco, CA)^67^. The heterodimeric RRLZ construct was generated by replacing the LZ domain with RRLZ, while the EELZ construct was created by stitching the KIF1A motor domain with EELZ and a HaloTag-StrepII tag. The K41 amber codon construct was generated by Q5 mutagenesis. All the constructs were validated by Sanger sequencing (Genomics Core Facility, Albert Einstein College of Medicine, Bronx, NY) or full plasmid sequencing (Azenta). The pAM87 plasmid containing a t-RNA synthetase and tRNA was a kind gift from Dr. Andreas Martin’s laboratory (University of California at Berkeley, Berkeley, CA)^68^. The Sfp (pET29b) plasmid was obtained from Addgene (#75015)^69^.

### E. coli-Based Protein Expression

All the constructs were expressed in *E. coli*. Each plasmid was transformed into BL21-CodonPlus(DE3)-RIPL competent cells (Agilent Technologies, #230280) and a single colony was picked and inoculated in 1 mL of terrific broth (TB)^70^ with 50 µg/mL carbenicillin and 50 µg/mL chloramphenicol. For Sfp expression, kanamycin was used instead of carbenicillin. The 1-mL culture was incubated at 37°C with shaking overnight and then transferred into 400 mL of TB containing 2 µg/mL carbenicillin and 2 µg/mL chloramphenicol. The culture was shaken at 37°C for 5 hours and then cooled to 16°C for 1 hour. Protein expression was induced by adding 0.1 mM IPTG, and the culture was incubated overnight at 16°C. Cells were harvested by centrifugation at 3,000×g for 10 minutes at 4°C, and the supernatant was discarded. The cell pellet was resuspended in 5 mL of B-PER™ Complete Bacterial Protein Extraction Reagent (ThermoFisher Scientific, #89821) supplemented with 2 mM MgCl_2_, 1 mM EGTA, 1 mM DTT, 0.1 mM ATP, and 2 mM PMSF. The suspension was flash-frozen in liquid nitrogen and stored at –80 °C until further use.

### E. coli-Based Expression of KIF1A K41amber

To express KIF1A with an amber codon, BL21-CodonPlus(DE3)-RIPL competent cells were first generated by transforming pAM-87 using the Mix & Go! *E. coli* transformation kit (Zymo Research, #T3001). The KIF1A K41amber plasmid was then transformed into the generated competent cells. The transformed cells were cultured in 1 L of terrific broth containing 25 µg/mL spectinomycin, 50 µg/mL carbenicillin, and 17 µg/mL chloramphenicol, following the same protocol as for other constructs. During the 1-hour cooling step prior to induction, 0.2 g of 4-azido-L-phenylalanine (AzF) (Vector Laboratories, #CCT-1406) was added to the culture. Induction with 0.1 mM IPTG and subsequent harvest steps were performed as described for other proteins.

### Purification of E. coli-Expressed Constructs

To purify *E. coli*-expressed protein, the frozen cell pellet was thawed at 37 °C and gently nutated at room temperature (RT) for 20 minutes to lyse the cells. For heterodimers, the two constructs were combined before the nutation step. For cysteine labeling, 1 mM TCEP was used in place of 1 mM DDT in all buffers. The lysate was cleared by centrifugation at 80,000 rpm (260,000×g, *k*-factor=28) for 10 minutes using a TLA-110 rotor in a Beckman Tabletop Optima TLX Ultracentrifuge. The resulting supernatant was passed through 500 μL of Ni-NTA Roche cOmplete™ His-Tag purification resin (Millipore Sigma, #5893682001) for His-tag tagged proteins. For heterodimers, the elute from Ni-NTA resin was subsequently passed through Strep-Tactin® 4Flow® high-capacity resin (IBA Lifesciences GmbH, #2-1250-010). The resin was washed with 10 mL of wash buffer (50 mM HEPES, pH 7.2, 300 mM KCl, 2 mM MgCl_2_, 1 mM EGTA, 1 mM DTT, 1 mM PMSF, 0.1 mM ATP, 0.1% (w/v) Pluronic F-127, 10% glycerol). Proteins were eluted with elution buffer containing 50 mM HEPES (pH7.2), 150 mM KCl, 2 mM MgCl_2_, 1 mM EGTA, 1 mM DTT, 1 mM PMSF, 0.1 mM ATP, 0.1% (w/v) Pluronic F-127, 10% glycerol. For His-tagged proteins, 150 mM imidazole was added to the elution buffer, while for Strep-II-tagged proteins, 5 mM desthiobiotin was included. For SNAPf-tag or Halo-Tag labeling, the appropriate ligand was added to the resin at a final concentration of 10 µM before elution and incubated at RT for 20 minutes. Excess ligand was removed by washing the resin with 5 mL of wash buffer. The final elute was flash-frozen in liquid nitrogen and stored at –80 °C.

### Purification of KIF1A K41F

The purification steps for the KIF1A K41F construct followed the same protocol as other proteins up to the wash step. After the resin was washed with 10 mL of wash buffer, an additional 5 mL of wash buffer without DTT was used to remove residual reducing agent. Subsequently, 1 mL of the wash buffer (without DTT) containing 150 µM 5,5-dithiobis(2-nitrobenzoic acid) (DTNB) was passed through the resin to protect surface cysteines. The resin was then washed with another 5 mL of wash buffer without DTT. For labeling, DBCO-LD655 (Lumidyne) was added to the resin to a final concentration of 0.1 mM, and the resin was incubated at 4 °C overnight in the dark. Following incubation, the resin was washed with 10 mL of wash buffer, and the protein was eluted following the same protocol as for other constructs.

### Microtubule Polymerization

To prepare stabilized MTs, 2 µL of 10 mg/mL tubulin (Cytoskeleton, #T240-B) was mixed with 2 µL of 1 mg/mL biotinylated tubulin (Cytoskeleton, #T333P-A), 1 µl of 1 mg/ml HiLyte488-labeled tubulin (Cytoskeleton, #TL488M-A), and 1 µL of 10 mM GTP. The mixture was incubated at 37°C for 20 minutes, after which 0.6 µL of 0.2 mM paclitaxel in DMSO was added. The incubation was continued for another 20 minutes. The resulting solution was carefully layered on top of 100 µL of glycerol cushion (80 mM PIPES, pH 6.8, 2 mM MgCl_2_, 1 mM EGTA, 60% (v/v) glycerol, 1 mM DTT, 10 µM paclitaxel) in a 230-µL TLA100 tube (Beckman Coulter, #343775), and centrifuged at 80,000×rpm (250,000×g, *k*-factor=10) for 5 minutes at RT in a Beckman Tabletop Optima TLX Ultracentrifuge. Following centrifugation, the supernatant was carefully removed, and the pellet was resuspended in 11 µL of BRB80G10 (80 mM PIPES, pH 6.8, 2 mM MgCl_2_, 1 mM EGTA, 10% (v/v) glycerol, 1 mM DTT, 10 µM paclitaxel). The MT solution was stored at RT in the dark until further use. For MT-binding and -release assays, only 5 µl of 10 mg/mL unlabeled tubulin was used for polymerization.

### Microtubule-Binding and -Release (MTBR) Assay

An MT-binding and -release (MTBR) assay was performed to remove inactive motors in preparation for single-molecule TIRF and MINFLUX assays. Protein samples (50–100 μL) of each construct were buffer-exchanged into a low-salt buffer (30 mM HEPES, pH 7.2, 50 mM KCl, 2 mM MgCl_2_, 1 mM EGTA, 1 mM DTT, 1 mM AMP-PNP, 10 µM paclitaxel, 0.1% (w/v) Pluronic F-127, and 10% glycerol) using 0.5-mL Zeba™ spin desalting column (7-kDa MWCO) (ThermoFisher Scientific, #89882). For ybbR-tag labeling, the protein solution was first mixed with Sfp (final 2 µM), 5 µM CoA-LD655, and 5 µM CoA and incubated at 4°C for 3 hours before buffer exchange. For cysteine labeling, maleimide-LD655 and maleimide-CF405 were added to the protein to final concertation of 0.1 mM, and the solution was incubated at 4 °C for 2 hours prior buffer exchange. After buffer exchange, the solution was warmed to RT, and 3 μL of 5 mg/mL paclitaxel-stabilized MTs was added. The solution was then thoroughly mixed and spun through a 100 μL glycerol cushion (80 mM PIPES, pH 6.8, 2 mM MgCl_2_, 1 mM EGTA, 1 mM DTT, 10 µM paclitaxel, and 60% glycerol) by centrifugation at 45,000 rpm (80,000×g, *k*-factor=33) for 10 minutes at RT in TLA-100 rotor using a Beckman Tabletop Optima TLX Ultracentrifuge. The supernatant was discarded, and the pellet was resuspended in 20–50 μL high-salt release buffer (30 mM HEPES, pH 7.2, 300 mM KCl, 2 mM MgCl_2_, 1 mM EGTA, 1 mM DTT, 10 μM paclitaxel, 1 mM ATP, 0.1% (w/v) Pluronic F-127, and 10% glycerol). The MTs were then removed by centrifugation at 40,000 rpm (60,000×g, *k*-factor=41) for 5 minutes at RT. Finally, the supernatant containing the active motors was aliquoted, flash-frozen in liquid nitrogen, and stored at –80 °C. For low-ATP assay, the supernatant was buffer-exchanged into a low-ATP buffer (30 mM HEPES, pH 7.2, 100 mM KCl, 2 mM MgCl_2_, 1 mM EGTA, 1 mM DTT, 10 μM paclitaxel, 10 µM ATP, 0.1% (w/v) Pluronic F-127, and 10% glycerol) prior to flash freezing.

## Total Internal Reflection Fluorescence (TIRF) Assay

### Flow Chamber Preparation

A flow chamber was assembled using a glass slide (Fisher Scientific, #12-550-123) and an ethanol-cleaned coverslip (Zeiss, #474030-9000-000) separated by two thin strips of parafilm to create a channel. The chamber was first treated with 10 µL of 0.5 mg/ml BSA-biotin (ThermoScientific, #29130), which was introduced into the chamber and incubated for 10 minutes. The chamber was then washed with 2×20 µL blocking buffer (80 mM PIPES, pH 6.8, 2 mM MgCl_2_, 1 mM EGTA, 10 µM paclitaxel, 1% (w/v) Pluronic F-127, 2 mg/mL BSA, 1 mg/mL α-casein) and incubated for 30 minutes to block non-specific surface interactions. Following the blocking step, 10 µL of 0.25 mg/ml streptavidin (Promega, #Z7041) was introduced into the chamber and incubated for 10 minutes. The chamber was washed again with 2×20 µL blocking buffer, after which 10 µL of 0.02 mg/ml biotin-labeled MTs in the blocking buffer was flowed into the chamber and incubated for 1 minute. The chamber was washed with a final 2×20 µL blocking buffer and stored in a humid chamber to prevent drying until further use.

### Sample Preparation

The motor solution was diluted to appropriate concentration, and 1 µL of the diluted motor solution was added to 50 µL of motility buffer (80 mM PIPES, pH 7.2, 2 mM MgCl_2_, 1 mM EGTA, 10 µM paclitaxel, 0.5% (w/v) Pluronic F-127, 5 mg/mL BSA, 1 mg/mL α-casein, 2 mM ATP, 2 mM biotin, gloxy oxygen scavenger system). A total of 2×20 µL of the mixture was flowed into the prepared flow chamber, and the chamber was sealed with vacuum grease to prevent evaporation. For experiments under nucleotide free apo conditions, 1 µL of apyrase (0.5 U/µl, NewEngland Biolabs, #M0398S) was added to the motility buffer instead of ATP. For ADP conditions, 1 µL of 100 mM ADP and 1 µL of 0.5 U/µL hexokinase were used in place of ATP.

### Data Acquisition

The images were acquired using BioVis software (BioVision Technologies) with an acquisition time of 200 ms per frame, capturing 600 frames per movie.

### Data Analysis

The kymographs were generated using Fiji^71^, and velocities were analyzed with a custom MATLAB program.

## MINFLUX Single-Molecule Assay

### Flow Chamber preparation

The flow chamber was assembled as described for the TIRF assay. After assemby, 10 µl of 150 nm gold colloid (BBI Solutions, #EM.GC150/4) was flowed into the chamber and incubated for 2 hours. The solution was then completely removed, and the chamber was kept at RT overnight. The subsequent steps were performed as described for TIRF assay.

### Sample Preparation

Sample preparation followed the same protocol as for the TIRF assay. For low ATP concentrations, phosphoenolpyruvate (PEP) and pyruvate kinase (PK) were added to the final solution to final concentrations of 1 mM and 0.1 mg/mL, respectively.

### Data Acquisition

MINFLUX tracking data were recorded using an Abberior Instruments MINFLUX microscope (Abberior Instruments, Göttingen, Germany) operated with iMSPECTOR Image Acquisition & Analysis Software v16.3.15636. Details of the microscope and its components were previously described^6^. The microscope is built around an inverted microscope body (IX83, Olympus) equipped with 405 nm, 485 nm, 561 nm, and 642 nm laser lines. Tracks were acquired using a 100x 1.45 NA UPLXAPO oil immersion objective (Olympus), with the confocal pinhole set to 0.8 AU. Signal detection was performed using two avalanche photodiodes (APDs) in the far-red spectral range (650 – 685 nm and 685 – 720 nm).

Tracking sequences were adjusted to prioritize spatial precision over temporal resolution, as detailed in Supplementary Table S1. The background threshold for identifying fluorophore signals over background noise was optimized for each sample. The initial laser power, before applying the power scaling, was set to 16 % (108 µW at the scanner) for tracking at 100 µM and 2mM ATP, and 8-10 % (54-68 µW at the scanner) for tracking at 10 µM ATP. For tracking at 10 µM ATP, the sequence was further modified to localize signals at a 1 kHz rate (scheduling parameter set to 1 ms), reducing photobleaching and accommodating the slower motor movement under these conditions.

### Data Analysis

The MINFLUX data were exported from iMSPECTOR as a *.mat* files and analyzed using custom MATLAB scripts. Initially, all traces (localizations with the same *tid* parameter) were evaluated based on the number of localizations, eccentricity (length-to-width ratio), and estimated tracking precision, which was calculated as:

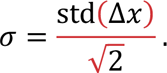

Here std() represents the standard deviation of the argument, and Δ*x* denotes the difference between neighboring localizations. Eccentricity was calculated as the maximal ratio of the eigenvalues of the covariance matrix of the *xy*-coordinates of each trace.

Traces with at least 500 localizations, an eccentricity greater 100, and a tracking precision below 6 nm in both *x* and *y* were further processed. First, the traces were binned to a time resolution of 1 ms, and the major axis of movement was aligned with the x-axis by projecting the *xy*-positions onto the eigenvector of the covariance matrix corresponding to the larger eigenvalue.

The stepwise motion along this aligned axis was analyzed using the MATLAB function *ischange*, with a threshold parameter proportional to *σ*^2^. The proportionality constant was determined as follows: for each ATP concentration, a large number of traces was fitted while varying the proportionality constant from 0 to 300. Steps smaller than 4 nm were considered indicative of overfitting, while steps larger 22 nm were categorized as overfitting. A threshold parameter of 100*σ*^2^ provided an optimal balance between minimizing underfitting and overfitting across all constructs and ATP concentrations. Based on the changepoints identified by *ischange*, the mean *xy*-positions and dwell times of each plateau were calculated. The processed data were saved and used to pool the results from multiple MINFLUX measurements.

### AlphaFold2 Prediction

The structures of kinesins were predicted using ColabFold (v1.5.5: AlphaFold2 using MMseqs2)^72^.

### Graphic Visualization

The molecular structures shown in the figures were visualized using UCSF ChimeraX^73^. The graphical models in Figures 5 and 7 were created using BioRender (https://www.biorender.com/).

**Figure 7.**
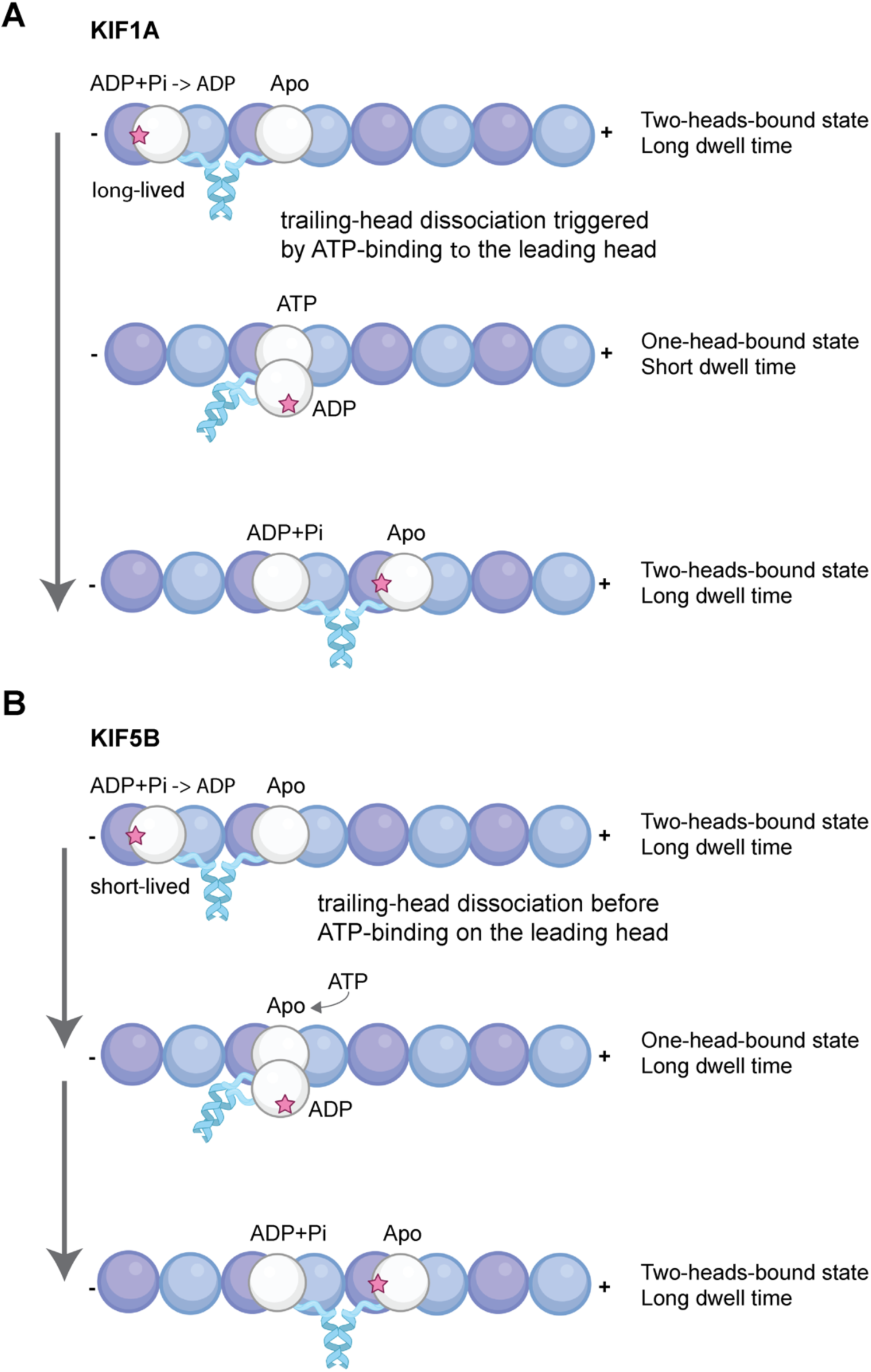
Proposed models for the stepping mechanisms of KIF1A and KIF5B. (**A**) Model for KIF1A stepping: The trailing head remains bound to the microtubules until the leading head binds ATP, resulting in a short-lived intermediate one-head-bound state. This mechanism ensures coordinated stepping and maintains high processivity. (**B**) Model for KIF5B stepping: The trailing head dissociates from the microtubule before the leading head binds to ATP, resulting in a prolonged one-head-bound state. This behavior contributes to reduced processivity compared to KIF1A.

## Acknowledgements

Lu. Rao and A. Gennerich thank Erik Jonsson for helpful discussions on unnatural amino acid (UAA)-based protein expression, Hernando Sosa for suggesting ybbR tag positions, Luke Lavis for providing JF dyes, and Wendy Chung for insightful discussions on the disease severity associated KIF1A mutations. UCSF ChimeraX, developed at the University of California, San Francisco, was supported by National Institutes of Health R01-GM129325 and the Office of Cyber Infrastructure and Computational Biology, National Institute of Allergy and Infectious Diseases. L. Rao and A. Gennerich were supported by National Institutes of Health grants 1R01GM147332.

## Author Contributions

L.R. generated, expressed, and purified constructs for MINFLUX- and TIRF-based single-molecule studies and prepared all samples. J.M. designed and conducted MINFLUX experiments, while L.R. designed and performed TIRF experiments. J.O.W. and L.R. developed the MATLAB program for MINFLUX data analysis. Data analysis and interpretation were carried out by J.O.W., L.R. and A.G. The research study was designed by L.R. and A.G. The manuscript was written by A.G, L.R. and J.O.W., with feedback from J.M. A.G. conceived the project, coordinated the research, and secured funding.

