## Supplementary Figures and Table for "A Two-Heads-Bound State Drives KIF1A Superprocessivity"

**Supplementary Files**

**Supplementary Tables**

**Table S1. Parameters for MINFLUX tracking sequences,** including iteration number (Itr), pattern diameter (L), target coordinate pattern (TCP), photon threshold, center-frequency-ration (CFR) limit, dwell time, pattern repeats within a given dwell time, background threshold, and power scaling factor. No CFR limit was applied during iterations 0-2.

| Itr | L (nm) | TCP | Photon threshold | CFR limit | Dell time (ms) | Pattern repeats | Background threshold (kHz) | Power factor |
|---|---|---|---|---|---|---|---|---|
| 0 | 284 | Hexagon | 40 | - | 0.4 | 1 | 70 | 1 |
| 1 | 302 | Hexagon | 20 | - | 0.4 | 1 | 70 | 1 |
| 2 | 151 | Hexagon | 10 | - | 0.4 | 1 | 40 | 2.0 |
| 3 | 76 | Hexagon | 10 | 0.9 | 0.4 | 1 | 40 | 2.5 |
| 4 | 29 | Hexagon | 10 | 2.0 | 0.1 - 0.3 | 1 | 130 - 250 | 6.0 |

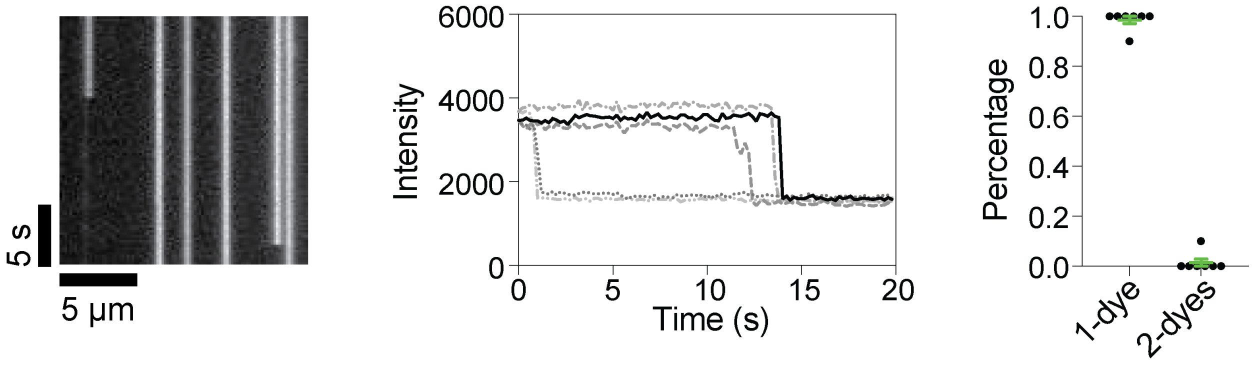


**Supplemental Figure 1. Photobleaching assay of LD655-labeled ybbR-tagged KIF1A. (A)** Representative kymograph showing the photobleaching of LD655-labeled KIF1A on a microtubule. **(B)** Example intensity traces from the photobleaching assay, illustrating discrete bleaching steps corresponding to individual fluorophores. **(C)** Dye counts on different microtubules. Microtubules (n=7); total molecules analyzed (n=53).

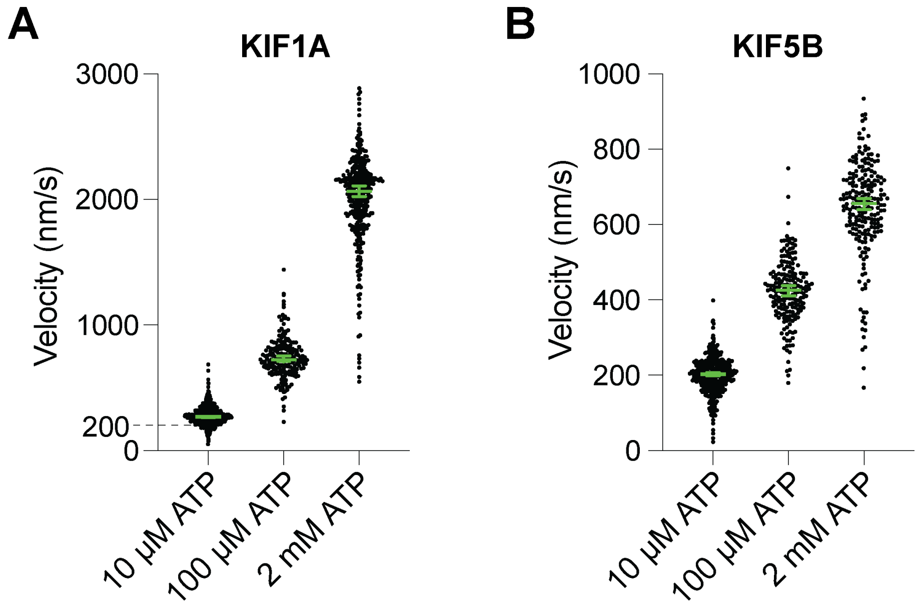


**Supplemental Figure 2. Velocity of KIF1A and KIF5B at different ATP concentration based on MINFLUX tracking (A)** Velocity of ybbR-tagged KIF1A at 10 μM (268 [263, 274] nm/s), 100 μM (723 [707, 752] nm/s), and 2 mM ATP (2063 [2025, 2108] nm/s). **(B)** Velocity of KIF5B S43C at 10 μM (203 [198, 207] nm/s), 100 μM (426 [411, 438] nm/s), and 2 mM ATP (656 [639 670] nm/s). Green bars indicate median values with 95% CIs.

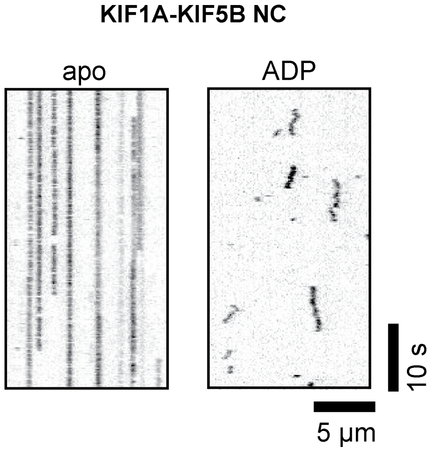


**Supplemental Figure 3. Kymograph examples of KIF1A-KIF5B NC chimera at apo and ADP states.** Representative kymographs showing the binding behavior of the KIF1A-KIF5B NC chimera on microtubules under apo conditions and in the presence of 1 mM ADP.

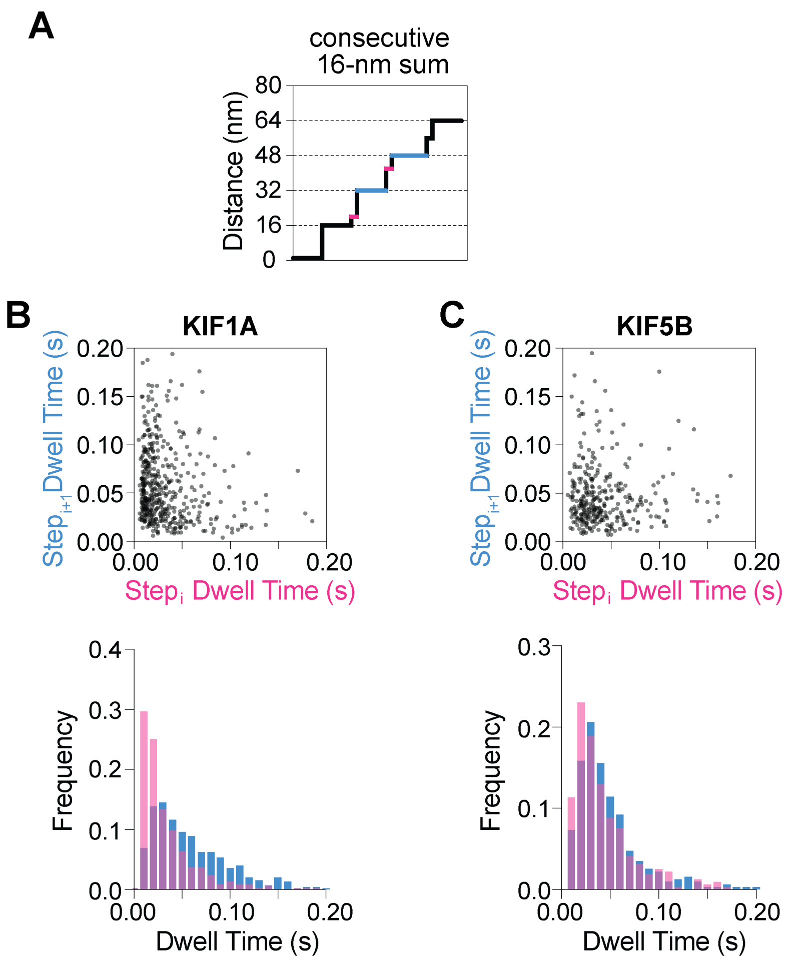


**Supplemental Figure 4. Dwell time analysis of sub-steps without restricting step sizes to 8 ± 2 nm. (A)** Illustrations of the dwell time analysis process, as described in Figure 5. **(B)** (Top) Correlation of sub-step dwell times for KIF1A: n=456, Pearson *r*=-0.1433 [-0.2320 -0.05213], two-tailed *P*=0.0022. (Bottom) Histograms of the A-sub-steps (red) and B-sub-steps (blue). **(C)** (Top) Correlation of sub-step dwell times for KIF5B: n=319, Pearson *r*=-0.04981 -0.1588, 0.06033], two-tailed *P*=0.3752. (Bottom) Histograms of the A-sub-steps (red) and B-sub-steps (blue).

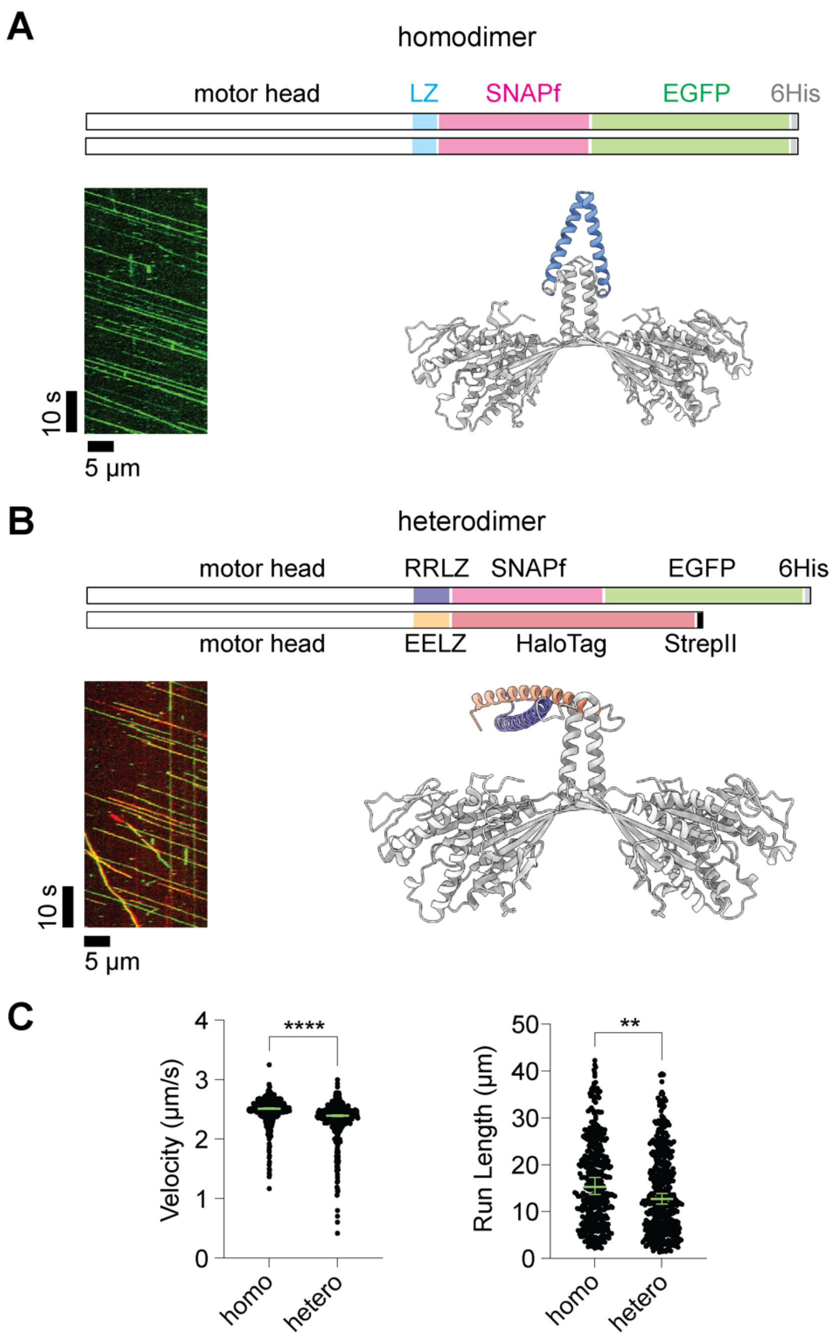


**Supplemental Figure 5. Heterodimeric KIF1A behaves similarly to homodimeric KIF1A. (A)** (Top) Schematic representation of the homodimeric KIF1A construct. (Bottom, left) Kymograph example of homodimeric KIF1A labeled with SNAP-TMR. (Bottom, right) ColabFold-predicted structure of homodimeric KIF1A with leucine zipper (LZ) (LZ, colored in blue). **(B)** (Top) Schematic representation of the heterodimeric KIF1A construct. (Bottom, left) Kymograph example of heterodimeric KIF1A labeled with SNAP-JF549 (green) and Halo-JF646[72] (red).

(Bottom, right) ColabFold-predicted structure of heterodimeric KIF1A with RRLZ-EELZ coiled coils. RRLZ is colored in purple, while EELZ is colored in orange. **(C)** Velocity and run length comparison between homodimeric and heterodimeric KIF1A. Velocity: Homodimer: 2.51 [2.50, 2.52] μm/s, n=419; heterodimer: 2.39 [2.38, 2.41] μm/s, n=440; **** two-tailed $P<0.0001$ (unpaired *t*-test with Welch's correction). Run length: homodimer: 15.26 [13.71, 17.26] μm; heterodimer: 12.70 [11.66, 13.84] μm; ** $P=0.0055$ (Kolmogorov-Smirnov test).
